# Loss of Purkinje cells in the developing cerebellum strengthens the cerebellothalamic synapses

**DOI:** 10.1101/2023.11.01.564864

**Authors:** Hiroshi Nishiyama, Naoko Nishiyama, Boris V. Zemelman

## Abstract

Cerebellar damage early in life often causes long-lasting motor, social, and cognitive impairments, suggesting the roles of the cerebellum in developing a broad spectrum of behaviors. This recent finding has promoted research on how cerebellar damage affects the development of the cerebral cortex, the brain region responsible for higher-order control of all behaviors. However, the cerebral cortex is not directly connected to the cerebellum. The thalamus is the direct postsynaptic target of the cerebellum, sending cerebellar outputs to the cerebral cortex. Despite its crucial position in cerebello-cerebral interaction, thalamic susceptibility to cerebellar damage remains largely unclear. Here, we studied the consequences of early cerebellar perturbation on thalamic development. Whole-cell patch-clamp recordings showed that the synaptic organization of the cerebellothlamic circuit is similar to that of the primary sensory thalamus, in which aberrant sensory activity alters synaptic circuit formation. The hemizygous deletion of the tuberous sclerosis complex-1 (*Tsc1*) gene in the Purkinje cell—known to cause Purkinje cell hypoactivity and autistic behaviors—did not alter cerebellothalamic synapses or intrinsic membrane properties of thalamic neurons. However, the ablation of Purkinje cells in the developing cerebellum strengthened the cerebellothalamic synapses and enhanced thalamic suprathreshold activities. These results suggest that the cerebellothalamic circuit is resistant to moderate perturbation in the developing cerebellum, such as the reduced firing rate of Purkinje cells, and that autistic behaviors are not necessarily linked to thalamic abnormality. Still, Purkinje cell loss alters the thalamic circuit, suggesting the vulnerability of the thalamus to substantial disturbance in the developing cerebellum.

## Introduction

A cerebellar abnormality is one of the most prominent characteristics of brain pathology in neurological disorders, such as autism spectrum disorders (ASDs), Down syndrome, schizophrenia, and spinocerebellar ataxia (Bailey *et al*., 1998; Pinter *et al*., 2001; Wang *et al*., 2014; D’Mello & Stoodley, 2015; Tamura *et al*., 2017; Moberget *et al*., 2017). The link between the cerebellum and cognitive impairments in these disorders previously seemed perplexing because the cerebellum was traditionally considered a part of the motor system. However, recent studies show that the cerebellum is also involved in various non-motor functions, such as cognitive processing, social behavior, fear, and emotion, through its interaction with other brain regions (Koziol *et al*., 2014; Palesi *et al*., 2015; Badura *et al*., 2018; Vaaga *et al*., 2020; Fastenrath *et al*., 2022; Chao *et al*., 2023).

Still, the emerging roles of the cerebellum in non-motor behaviors do not fully explain how the cerebellum contributes to the pathogenesis of the above neurological disorders. While cerebellar injury at birth is one of the highest risk factors for ASDs, damage in the mature cerebellum is not (Wang *et al*., 2014). Chemogenetic perturbation of cerebellar circuits causes severe social and cognitive impairments when it is done in the juvenile period but not in adulthood (Badura *et al*., 2018). These findings show that early cerebellar damage or dysfunction causes more profound neurological symptoms than later ones, suggesting that the cerebellum is necessary to develop functionally precise brain circuits.

Neuronal activity plays a crucial role in the maturation and refinement of synaptic circuits (Katz & Shatz, 1996; Lichtman & Colman, 2000). Therefore, aberrant activities of the immature cerebellum due to injury or genetic mutation potentially interfere with the development of its downstream targets (Wang *et al*., 2014). In this regard, recent studies focused on the cerebral cortex, the higher-order center for all behaviors, showing that early cerebellar damage in human patients results in reduced cortical volume (Limperopoulos *et al*., 2010, 2014; Stoodley & Limperopoulos, 2016). However, the cerebellum is not directly connected to the cerebral cortex. How aberrant cerebellar activities impair cortical development is largely unclear.

We focused on the thalamus in this study for the following reasons. First, the cerebellum sends dense, mono-synaptic excitatory projections to the thalamus, directly impacting the synaptic activities in the thalamus (Evarts & Thach, 1969; Chan-Palay, 1977; Bosch-Bouju *et al*., 2013). Second, the thalamus is the heart of cerebello-cerebral interaction. It relays cerebellar activity to the motor and non-motor areas of the cerebral cortex (D’Angelo & Casali, 2013; Koziol *et al*., 2014); hence, if thalamic circuits are not formed properly, functional interaction between the cerebellum and cerebral cortex must be disturbed. Third, neuronal activities are necessary for the maturation of the primary sensory thalamus (Hooks & Chen, 2006; Wang & Zhang, 2008; Takeuchi *et al*., 2014), suggesting that normal cerebellar activities are also required to build functional cerebellothalamic circuits.

To study how cerebellar perturbation affects thalamic development, we compared neuronal function in two mouse lines. We created a transgenic mouse line in which cerebellar Purkinje cells are ablated from the first postnatal week. This mouse line represents a considerable perturbation because the ablation of Purkinje cells removes their potent tonic inhibition on the deep cerebellar nuclei, substantially altering the cerebellar output activities to its downstream targets. We also used a previously reported mouse model of autism caused by Purkinje cell-specific hemizygous deletion of the *Tsc1* gene from the first postnatal week to study the effect of a moderate perturbation (Tsai *et al*., 2012).

We performed whole-cell patch-clamp recordings in thalamic slices obtained from these mice and characterized the synaptic and intrinsic properties of the cerebellothalamic circuit. While the hemizygous deletion of the *Tsc1* gene did not cause any significant changes, Purkinje cell ablation increased the number of synaptically-evoked thalamic action potentials, likely due to strengthened cerebellothalamic synapses. These results show that damage in the developing cerebellum potentially alters the cerebellothalamic circuit, depending on the type and severity of the damage.

## Materials and Methods

### Animals

All animal procedures were performed in accordance with the University of Texas at Austin Institutional Animal Care and Use Committee guidelines.

The knock-in mouse line, in which the Cre-inducible DTA gene is inserted downstream of the neuron-specific enolase promoter (*Eno2^fsDTA/+^*; RBRC10744, Riken BioResource Center, Japan), the transgenic mouse line, in which Purkinje cells-specific protein-2 promoter drives Cre expression (*Pcp2^Cre/+^*; JAX:004146, The Jackson Laboratory), and the floxed mutant mouse line, in which exons 17 and 18 of the *Tsc1* gene are flanked by loxP sites (*Tsc1^flox/+^*; JAX:005680, The Jackson Laboratory) were used in this study. These mouse lines were maintained in the C57BL/6J background.

To ablate Purkinje cells, we crossed *Pcp2^Cre/+^* mice with *Eno2^fsDTA/+^* mice and obtained the double transgenic mice (*Pcp2^Cre/+^ Eno2^fsDTA/+^*). The single transgenic littermates (either *Pcp2^Cre/+^* or *Eno2^fsDTA/+^*) were used as a control group. To delete the *Tsc1* gene in Purkinje cells, we crossed *Pcp2^Cre/+^* mice with *Tsc1^flox/+^* mice and obtained the double transgenic mice (*Pcp2^Cre/+^ Tsc1^flox/+^*). The single transgenic littermates (either *Pcp2^Cre/+^* or *Tsc1^flox/+^*) were used as a control group.

### Virus preparation

An adeno-associated virus (AAV) encoding an enhanced human synapsin promoter, channelrhodopsin2-tdTomato, woodchuck hepatitis virus posttranscriptional regulatory element (WPRE) and SV40 polyadenylation signal was assembled using a modified helper-free system (Stratagene) as a serotype 2/1 (rep/cap genes) AAV and harvested and purified over sequential cesium chloride gradients as previously described (Grieger *et al*., 2006). The hybrid synapsin promoter comprised a cytomegalovirus enhancer (Niwa *et al*., 1991), a neural-restrictive silencer element (Mori *et al*., 1992) and a human synapsin promoter (Kügler *et al*., 2001). The channerhodopsin2 fusion protein included C-terminal Golgi and endoplasmic reticulum export signals (Ma *et al*., 2001; Hofherr *et al*., 2005) to aid membrane expression. Virus titers were typically >10^13 genomes per ml.

### Animal injection

The injection was performed on neonatal male and female mice around the postnatal day 7 (P7) before substantial cerebellar atrophy occurred in the Purkinje cell ablation mice. AAV was injected into the deep cerebellar nuclei of the control mice, Purkinje cell-specific *Tsc1* deletion mice, and Purkinje cell ablation mice.

Neonatal mice were anesthetized with 5% isoflurane, and the dose was reduced to 1.5-2% following the induction. Carprofen (10 mg/kg) was subcutaneously administrated as an analgesic. The scalp was cut, and the left side of the interparietal bone was exposed. Lidocaine (2%) was topically applied to the exposed skull and adjacent muscle surfaces. Posterior to the lambdoid suture, another suture runs mediolaterally between the interparietal and occipital bones. We used the midpoint of this suture as a landmark and located the injection site 1.5 mm left and 1.2 mm anterior from the landmark. A small craniotomy was made on the injection site with a dental drill and a drill bit (0.8 mm tip diameter), and a glass pipette (20-40 μm tip diameter) containing the virus solution was inserted vertically to the depth of 2.2 mm from the surface. Approximately 200-300 nl of virus was injected over 4-6 minutes using a Nanoject II injector (Drummond Scientific). This injection coordinates primarily label the interposed nucleus of the deep cerebellar nuclei, but the virus also spreads into the lateral and medial nucleus. The pipette was left in place for 5 minutes before being withdrawn, and the scalp was sutured. The mice were returned to their mother after recovering from anesthesia. Carprofen (10 mg/kg) was administrated every 24 hours for 2 days post-surgery as an analgesic. The mice were kept alive for 1.5-3 months until used for electrophysiological recordings.

### Histochemistry

The control and Purkinje cell ablation mice (1-month-old) were anesthetized with an intraperitoneal injection of ketamine/xylazine (100/10 mg/kg) and intracardially perfused with 4 % paraformaldehyde in phosphate-buffered saline (PBS). The brains were extracted post-perfusion and immersed in the same fixative at 4 °C overnight. After washing the tissues with PBS, sagittal sections of the cerebellum (50 µm thickness) and coronal sections of the forebrain (80 µm thickness) were cut using Microm HM650V vibration microtome (Thermo Fisher Scientific). The sections were washed with PBS containing 0.3 % Triton X-100 (PBST).

Then, the cerebellar sections were blocked at room temperature for at least one hour in the blocking solution containing PBST and 5 % normal goat serum. Following the blocking step, the sections were incubated with anti-calbindin-D28k mouse monoclonal antibody (1:2000 dilution by the blocking solution, C9848, Millipore Sigma) at 4 °C overnight, then washed with PBST and incubated with goat anti-mouse IgG (H+L) Alexa Fluor 594 (1:500 dilution by the blocking solution, A-11032, Thermo Fisher Scientific) at room temperature for at least 2 hours. Sections were then washed with PBST and mounted with 4’, 6-diamidino-2-phenylindole (DAPI)-containing mounting media (DAPI Fluoromount-G, 0100-20, Southern Biotech).

The forebrain sections were incubated with Blue NeuroTrace Fluorescent Nissl Stains (1:100 dilution by PBS, N-21479, Thermo Fisher Scientific) at room temperature for 2-3 hours. Sections were then washed with PBST and mounted with mounting media (Fluoromount-G, 0100-01, Southern Biotech).

### Image acquisition

The wide-field and confocal fluorescent images were acquired with an Olympus FV1000 confocal microscope system (Olympus, Japan). For the wide-field images, the stained sections were excited by a mercury arc lamp, and the fluorescent emission was collected by a 4× objective lens (0.1 NA). DAPI and Blue NeuroTrace Fluorescent Nissl were excited using a bandpass filter 325-375 nm (violet), and the fluorescent emission was filtered with a bandpass filter 435-485 nm (blue) before being captured by a CCD camera. A long-pass filter 400 nm was used as a dichroic mirror to separate the violet excitation and blue emission. Alexa Fluor 594 was excited using a bandpass filter 510-560 nm (green), and the fluorescent emission was filtered with a bandpass filter 575-645 nm (red). A long-pass filter 565 nm was used as a dichroic mirror to separate the green excitation and red emission.

For the confocal images, the immunostained cerebellar sections were excited by 405 nm (for DAPI) and 543 nm (for Alexa Fluor 594) lasers. The DAPI and Alexa Fluor 594 fluorescent emissions were separated by a long-pass filter 490 nm and then filtered by bandpass filters 430– 470 nm and 560–660 nm, respectively. Z-stack images were acquired with a 40× objective lens (0.75 NA) at 2 µm step size with the x-y resolution of 0.8 µm/pixel.

### Electrophysiology: overall procedures

Acute brain slices were cut near physiological temperatures, as a previous study recommended (Huang & Uusisaari, 2013). The cutting and holding solutions were prepared by referring to the methods optimized for adult brains (Ting *et al*., 2014, 2018), although we used sucrose to replace NaCl and simplified the procedure.

Mice (1.5-3 months old, 5-11 weeks post-injection) were deeply anesthetized with isoflurane. The cerebellum was quickly removed after decapitation and submerged in a warmed cutting solution (32-34 ℃) containing (in mM) 150 Sucrose, 30 NaHCO_3_, 20 HEPES, 2.5 KCl, 1.2 NaH_2_PO_4_, 0.5 CaCl_2_, 10 MgCl_2_, 5 Na-ascorbate, 3 Na-Pyruvate, and 25 dextrose. Coronal forebrain slices (220-µm-thick) containing the ventrolateral thalamus (VL) were cut in the solution using a 7000smz-2 vibration microtome (Campden Instruments), recovered in a warmed holding solution (32-34 ℃) containing (in mM) 90 NaCl, 20 Sucrose, 30 NaHCO_3_, 20 HEPES, 2.5 KCl, 1.2 NaH_2_PO_4_, 2 CaCl_2_, 2 MgCl_2_, 5 Na-ascorbate, 3 Na-Pyruvate, and 25 dextrose. The slices were kept at the warmed temperature for 30 minutes and then stored at room temperature until used for recording.

Recordings were performed at 32 ℃ in a submerged chamber perfused with artificial cerebrospinal fluid (ACSF) containing (in mM) 124 NaCl, 2.5 KCl, 1.25, NaH_2_PO_4_, 25 NaHCO_3_, 2 CaCl_2,_ 2 MgCl_2_, 20 dextrose, supplemented with Gabazine (5 μM, HB0901, Hello Bio) to block GABA_A_ receptor-mediated inhibitory synaptic transmission. To block α-amino-3-hydroxy-5-methyl-4-isoxazolepropionic acid receptors (AMPA-R) and N-methyl D-aspartate receptors (NMDA-R)-mediated excitatory synaptic transmission, the AMPA-R antagonist 2,3-Dioxo-6-nitro-1,2,3,4-tetrahydrobenzo[*f*]quinoxaline-7-sulfonamide (NBQX;10 μM HB0443, Hello Bio) and NMDA-R antagonist DL-2-Amino-5-phosphonopentanoic acid (DL-APV; 100 μM HB0252, Hello Bio) were added to ACSF. All the solutions were bubbled with 95% O_2_ and 5% CO_2_, and the osmolarity was adjusted around 310 ± 10 mOsm/kg.

Whole-cell patch-clamp recordings were made using a Multiclamp 700B amplifier (Molecular Devices) and AxoGraph data acquisition software (AxoGraph). The recording data were sampled at 20 kHz with an ITC-18 AD/DA board (Heka). The intracellular solution for voltage-clamp recordings contained (in mM) 35 CsF, 100 CsCl, 10 EGTA, 10 HEPES, 4 QX314 (∼290 mOsm/kg, pH adjusted to ∼7.3 with CsOH). The intracellular solution for current-clamp recordings contained (in mM) 130 K-MeSO_3_, 4 KCl, 10 HEPES, 8 Na_2_-phosphocreatine, 0.2 EGTA, 4 Mg-ATP, 0.4 Na-GTP (∼290 mOsm/kg, pH adjusted to ∼7.3 with KOH). The pipette resistance was around 3 ± 0.5 MΩ when filled with the intracellular solutions.

Cerebellothalamic axon terminals expressing ChR2 were identified in the VL using tdTomato fluorescence excited by 560 nm wavelength LED light (BLS-LCS-0560-03-22, Mightex). Individual postsynaptic VL neurons were visualized with a Nikon Fluor 60X water immersion objective lens (1.0 NA) under differential interference contrast mode. After whole-cell recording was established, ChR2 was excited by a brief (0.1-0.2 msec) pulse of 470 nm wavelength LED light (BLS-LCS-0470-03-22, Mightex). The duration, power, and timing of the pulse were controlled by BLS-series software (BLS-SA02-US, Mightex). The power under the objective lens was measured using an optical power meter (PM10, Thorlabs) and a sensor (D10MM, Thorlabs). The synaptic and intrinsic properties of VL neurons were quantified as described below. The AxoGraph recording data files were read with Python using the Axographio package, and all the measurements, threshold detection, and curve fitting were performed with Python.

### Voltage-clamp recordings

VL neurons were voltage-clamped at -70 mV, and ChR2-expressing cerebellothalamic axons were stimulated every 20 seconds by a 470 nm light pulse. The light power density was gradually increased from 0 to ∼40 mW/mm^2^. This stimulation protocol yielded one or several discrete steps of excitatory postsynaptic currents (EPSCs). The peak amplitude and charge transfer of each EPSC step were quantified and plotted against the light density. The charge transfer vs. light density plots and the corresponding EPSC traces were used to count the number of discrete EPSC steps.

After the above recording session, the same VL neurons remained held at -70 mV and stimulated by a pair of light pulses with the maximum power. The paired light pulses were given every 20 seconds by changing the paired-pulse interval from 25 to 200 msec. This recording was repeated 2-4 times, and the EPSC traces for each paired-pulse interval were averaged. Finally, the holding potential was changed from -70 mV to +40 mV, and the same neurons were stimulated by the maximum power of light. The light pulse was given every 20 seconds 3-6 times, and the EPSC traces were averaged.

The amplitude, 10-90 % rise time, and the weighted decay time constant of the first EPSC in the paired-pulse recording were measured. The weighted decay time constant was calculated as described in a previous study (Cathala *et al*., 2000). Briefly, a double exponential decay curve was fitted to the decay phase of the EPSCs. If the coefficient and time constant of a fast and slow decay component are C_fast_, C_slow_, τ_fast_, and τ_slow_, the weighted decay constant was calculated as: (C_fast_τ_fast_ + C_slow_τ_slow_) / (C_fast_ + C_slow_). The paired-pulse ratio was calculated as the amplitude ratio between the second and the first EPSCs. The AMPA/NMDA ratio was calculated as the amplitude ratio between the EPSC at -70 mV and the EPSC + 40 mV.

The series resistance was measured in all recordings by the current response to a 5 mV, 50 msec hyperpolarizing voltage step. The cells showing a series resistance larger than 15 MΩ were excluded. In addition, some cells showed no or very small EPSCs, most likely due to their primary synaptic inputs being damaged or not labeled by ChR2. To reduce the contribution of those cells to underestimating the synaptic strength and the number of synaptic inputs, we excluded the cells showing their maximum EPSC amplitude below 1 nA.

Furthermore, a few VL neurons expressed ChR2. These neurons were directly stimulated, showing fundamentally different currents from EPSCs, and were excluded from analysis. The VL neurons may have been infected following virus diffusion through the superior cerebellar peduncle to the thalamus. While trans-synaptic labeling has previously been postulated (Castle *et al*., 2014; Zingg *et al*., 2017, 2020), we have seen no evidence of AAV transfer between cells with any of our reagents.

### Current-clamp recordings

VL neurons were initially voltage-clamped at -70 mV and stimulated with light. Cells were discarded if they were directly stimulated or showed no or small EPSCs. Other cells were switched to current-clamp mode with no holding current. The series resistance was compensated using bridge balance.

Cells were stimulated every 20 seconds with a single light pulse, and the number of action potentials (APs) evoked by the maximum power of light was counted. The average voltage before the light pulse was used to calculate the resting membrane potential. The input resistance was calculated from the average steady-state voltage deflection in response to a 50 pA, 300 msec hyperpolarizing current injection.

The synaptically evoked APs began to fire on the steep rise of excitatory postsynaptic potentials, making the detection of the first AP threshold unreliable. Therefore, after the above recording session, we injected depolarizing current steps (200 msec, 25-250 pA with 25 pA increment) into the same cells to quantify AP properties. The AP threshold was defined as the voltage where dV/dt exceeded 10 mV/msec. The AP amplitude was defined as the difference between the peak and threshold. The AP half-width was defined as the duration during which an AP was higher than half of its amplitude. These values were obtained for the first AP in each current step and averaged across all steps that evoked APs.

In addition, the average voltage before the current injection was used to calculate the resting membrane potential. This value was averaged with the corresponding value obtained in the optogenetic stimulation and used as the resting membrane potential of the cells.

### Statistical analysis

Bootstrapping was used for most statistical analyses. This method does not require the sample or population to be normally distributed. The R package, Dabestr (Ho *et al*., 2019), was used to resample the observed data 5,000 times. It computes the observed mean difference between the control and cerebellar mutant mice, the resampled mean difference, and its distribution and 95% confidence interval. If the 95% confidence interval of the resampled mean difference does not include zero, it is equivalent to *p* < 0.05 in null hypothesis significance testing. Furthermore, beyond the statistical significance, the sharpness of the resampled distribution and the proximity of the confidence interval to zero allows for inferring the certainty of the difference (Calin-Jageman & Cumming, 2019; Ho *et al*., 2019).

For the paired-pulse ratio, mixed model analysis of variance (Mixed ANOVA) was used to test the difference between the control and cerebellar mutant mice, the interstimulus intervals, and the interaction between these variables. The *p*-value less than 0.05 was considered statistically significant.

For the correlation, Pearson’s correlation coefficient was computed between the two variables, and the *p*-value less than 0.05 was considered a statistically significant correlation. All the statistical analyses and data visualization were performed with R.

## Results

While the primary goal of this study is to gain basic biological insights into how the cerebellum contributes to thalamic formation, we also sought to understand it in the context of neurological disorders. Therefore, we perturbed the developing cerebellum in a manner that resembles human diseases. One of the most common cerebellar pathologies in neurological conditions is Purkinje cell abnormality (Gill & Sillitoe, 2019; Cook *et al*., 2021). Purkinje cells spontaneously discharge action potentials and tonically inhibit the deep cerebellar nuclei, the sole output of the cerebellum (Bell & Grimm, 1969; Latham & Paul, 1971; Palay & Chan-Palay, 1974). Hence, Purkinje cell dysfunction directly impacts the activity of the deep cerebellar nuclei, altering the communication between the cerebellum and other brain regions.

Loss of Purkinje cells or reduced firing rate is commonly associated with various neurological disorders, such as ASD, schizophrenia, bipolar disorder, Huntington’s Disease, and spinocerebellar ataxia (Zoghbi & Orr, 1995; Bailey *et al*., 1998; Kordasiewicz & Gomez, 2007; Maloku *et al*., 2010; Dougherty *et al*., 2012, 2013; Cook *et al*., 2021). To highlight the effects of Purkinje cell loss, we ablated Purkinje cells selectively by diphtheria toxin fragment A (DTA). A knock-in mouse in which the Cre-inducible DTA gene is inserted downstream of the neuron-specific enolase promoter (*Eno2^fsDTA/+^* mouse) was crossed with a transgenic mouse in which Purkinje cells-specific protein-2 promoter drives Cre expression (*Pcp2^Cre/+^* mouse). Since DTA expression is Cre-dependent, only Cre-expressing cells are ablated by DTA.

Previous studies used the *Eno2^fsDTA/+^* mouse successfully for cell type-specific ablation (Kobayakawa *et al*., 2007; Imayoshi *et al*., 2008; Kobayashi *et al*., 2013). The *Pcp2^Cre/+^* mouse is a well-established genetic tool for Purkinje cell-specific gene manipulation, in which Cre-mediated recombination begins from postnatal day 6 (P6) (Barski *et al*., 2000; Tsai *et al*., 2012; Sługocka *et al*., 2017). Single transgenic mice carrying either the DTA or the Cre gene were indistinguishable from wild-type mice (Fig. 1A-D). On the other hand, the double-transgenic mice expressing DTA in Purkinje cells showed cerebellar atrophy due to substantial loss of Purkinje cells (Fig. 1E-H). The atrophy was specific to the cerebellum; the overall forebrain structure was unaffected (Fig. 1B and F).

**Figure 1.**
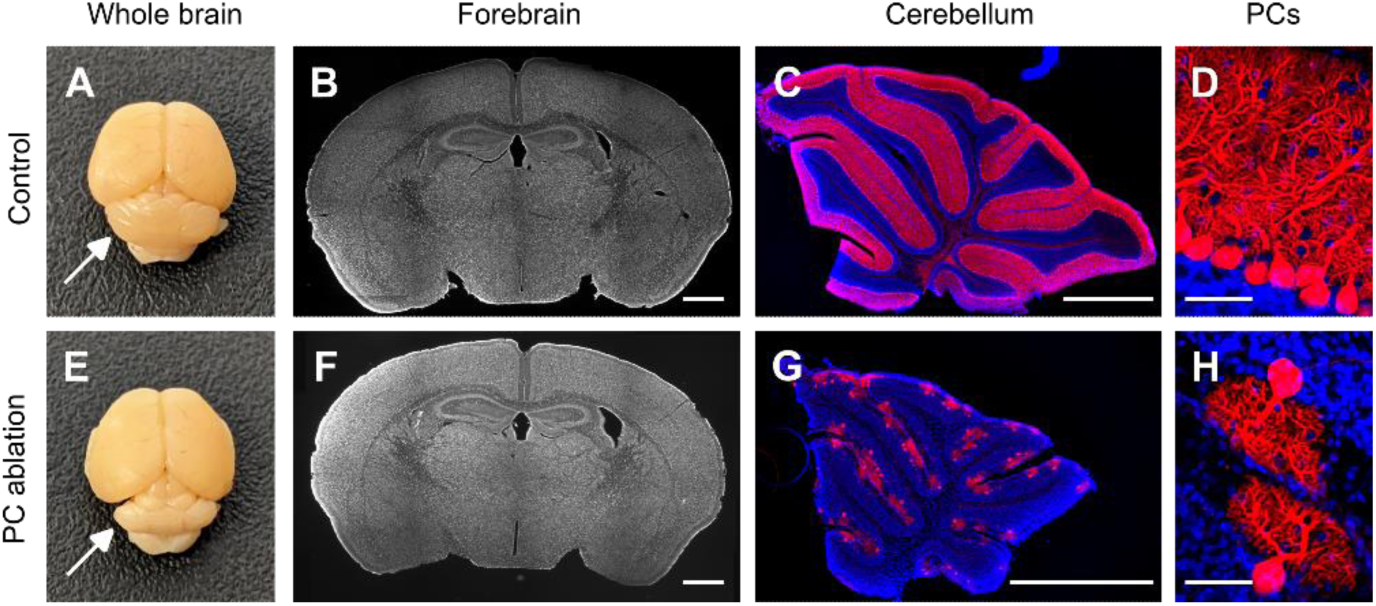
Selective ablation of Purkinje cells (PCs). **(A-D)** A control mouse carrying Cre-inducible DTA gene (*Eno2^fsDTA/+^*). This single transgenic mouse does not encode Cre; thus, DTA is not expressed. **(E-H)** A double transgenic mouse carrying Cre and DTA genes (*Pcp2^Cre/+^ Eno2^fsDTA/+^*). The PC-specific promoter Pcp2 drives Cre expression in this mouse. Therefore, DTA is expressed only in the PCs and ablates them. **(A and E)** The whole brain. The arrows indicate the cerebellum. Note that the cerebellum of the PC ablation mouse is substantially smaller than the control mouse. **(B and F)** Fluorescence Nissl stain of the forebrain, taken by a wide-field microscope. Scale bars: 1 mm. **(C and G)** Immunostaining of PCs with antibodies against a PC marker, calbindin-D28k (red), counterstained with DAPI (blue). The images were taken by a wide-field microscope. Scale bars: 0.5 mm. **(D and H)** Higher magnification of (C) and (G), taken by a confocal microscope. Scale bars: 50 μm.

While the Purkinje cell ablation mouse is a powerful model for studying the cerebellar contribution to thalamic development, the degree of cell loss exceeds the conditions found in most neurological disorders. To perturb the developing cerebellum in a manner that more closely mimics human diseases, we also used the *Pcp2^Cre/+^* mouse to delete the Tuberous sclerosis complex-1 (*Tsc1*) gene in Purkinje cells. The Purkinje cell-specific *Tsc1* deletion mouse is an established mouse model for autism, and administration of rapamycin from P7 to P35 rescues their impaired social behaviors (Tsai *et al*., 2012; Gibson *et al*., 2022). It suggests that TSC1 in Purkinje cells is crucial for a developmental period, making this mouse model particularly suitable for this study.

### Organization of cerebellothalamic synapses

The ventrolateral thalamus (VL) is a part of the motor thalamus, but the organization of long-range polysynaptic loops, i.e., cerebellum → thalamus → neocortex → pons → cerebellum, is essentially the same regardless of whether the cerebellum connects to the motor or non-motor areas of the neocortex (Kelly & Strick, 2003; D’Angelo & Casali, 2013; Koziol *et al*., 2014). To characterize the physiological properties of VL cerebellothalamic synapses, we injected adeno-associated virus (AAV) expressing channelrhodopsin-2-tdTomato (ChR2) into the deep cerebellar nuclei (Fig. 2A). The ChR2-expressing cerebellothalamic axons were optogenetically stimulated, and whole-cell voltage-clamp recordings of VL neurons were performed (Fig. 2B).

**Figure 2.**
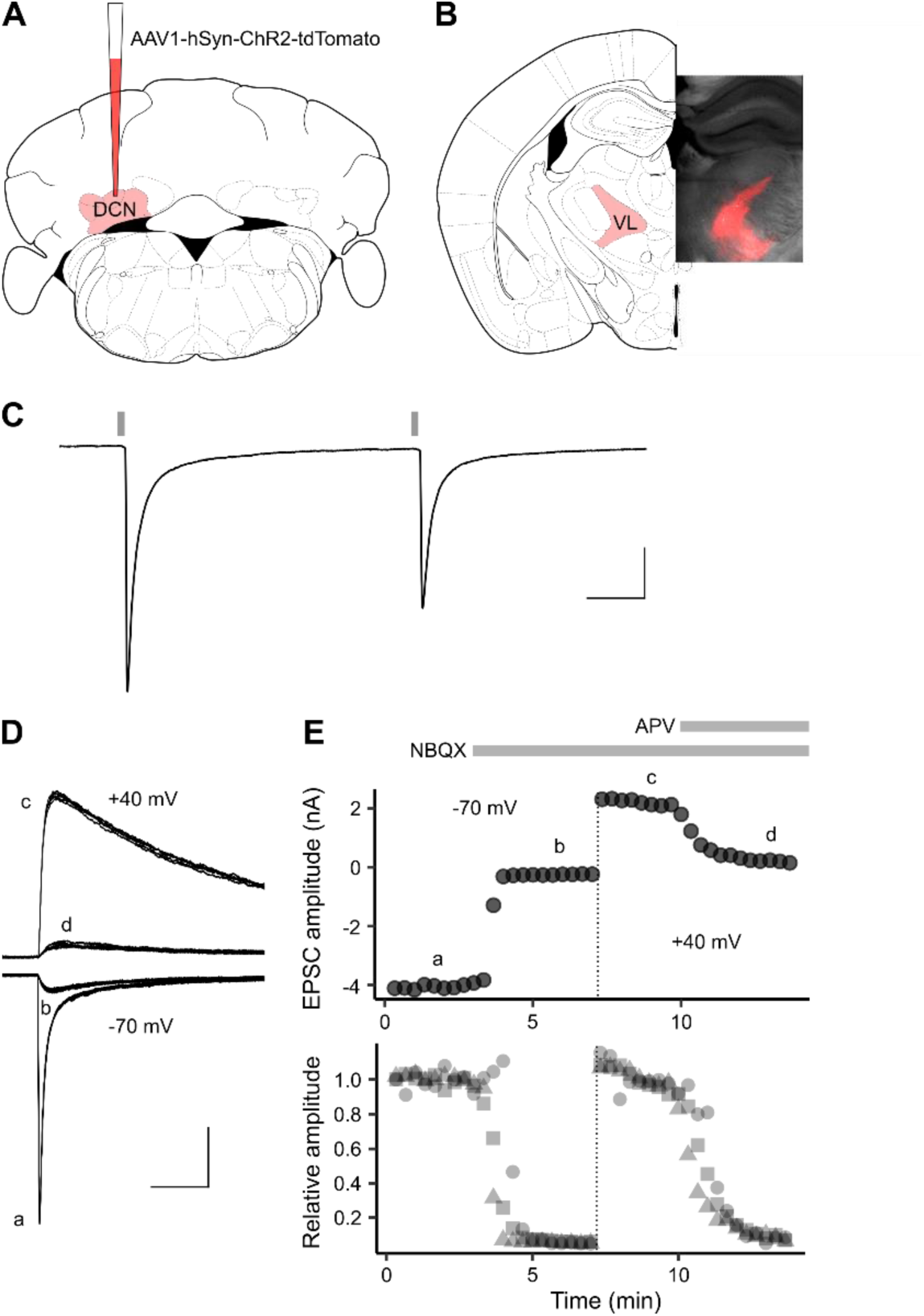
Recording of cerebellothalamic synapses. **(A)** AAV1 encoding ChR2-tdTomato under the control of the human synapsin promoter (hSyn) was injected into the deep cerebellar nuclei (DCN). **(B)** The ChR2-tdTomato expressing cerebellothalamic axons (red) in the ventrolateral thalamus (VL), shown with the corresponding region of the mouse brain atlas. **(C)** Whole-cell voltage-clamp recording from a VL neuron at -70 mV. A pair of light pulses (gray rectangles) evoked large EPSCs, showing paired-pulse depression. Scale bars: 1 nA and 20 msec. **(D)** EPSCs in another VL neuron. The holding potential was initially -70 mV (bottom), then switched to + 40 mV (top). AMPA receptor antagonist NBQX (10 μM) and NMDA receptor antagonist DL-APV (100 μM) were perfused during the recording. Scale bars: 1 nA and 20 msec. **(E)** The time course of the recording with antagonist application (gray bars). The vertical dashed lines indicate the timing of the holding potential switch. The top panel represents the neuron shown in panel D. The marks “a,” “b,” “c,” and “d” indicate the timing when the corresponding EPSCs in panel D were recorded. The bottom panel combines the recordings from three cells. The EPSC amplitude was normalized to the values before the NBQX application (-70 mV) and DL-APV application (+ 40 mV) in each neuron, represented by a different marker. All recordings in this figure were performed using control mice.

Consistent with the previous studies, optogenetic stimulation of cerebellothalamic axons evoked a large excitatory postsynaptic current (EPSC), showing paired-pulse depression (Fig. 2C) (Gornati *et al*., 2018; Schäfer *et al*., 2021). This type of EPSC is characteristic of driver inputs to the primary sensory thalamus, such as retinal inputs to the lateral geniculate nucleus and somatosensory inputs to the ventral posteromedial nucleus (Reichova & Sherman, 2004; Sherman, 2005; Bosch-Bouju *et al*., 2013). These sensory driver inputs evoke EPSCs primarily through AMPA- and NMDA-type glutamate receptors, and the ratio of the EPSCs recorded at -70 mV vs. at +40 mV is routinely used to measure the developmental increase of the AMPA/NMDA ratio (Chen & Regehr, 2000; Arsenault & Zhang, 2006; Hooks & Chen, 2006; Wang & Zhang, 2008; Takeuchi *et al*., 2014).

To test whether the same approach can be used in the cerebellothalamic synapses, we characterized the EPSC components at -70 mV and +40 mV in the VL of the control mice (Fig. 2D and E). The AMPA-R antagonist NBQX (10 μM) blocked 94.0 ± 0.6 % (mean ± standard deviation, n = 3 cells) of EPSCs at -70 mV, and the NMDA-R antagonist DL-APV (100 μM) blocked 92.1 ± 1.2 % (n = 3 cells) of EPSCs at +40 mV. Although there was a small remaining current at both holding potentials, these data indicate that the EPSC ratio recorded at -70 mV vs. +40 mV in the VL can be used to measure the AMPA/NMDA ratio like the primary sensory thalamus.

As the AMPA/NMDA ratio increases during the maturation of the primary sensory thalamus, neuronal activity-dependent synaptic pruning occurs. In the primary visual and somatosensory thalamus, individual thalamic neurons are initially innervated by many sensory driver inputs, but they are subsequently pruned during the first few postnatal weeks (Chen & Regehr, 2000; Arsenault & Zhang, 2006; Hooks & Chen, 2006; Wang & Zhang, 2008; Takeuchi *et al*., 2014). Consequently, a mature neuron in these thalamic nuclei receives only a few sensory driver inputs. To examine whether the mature VL neurons exhibit a similar innervation pattern, we recorded AMPA currents (AMAP-EPSCs) in the control mice by gradually increasing the power density of the blue light (Fig. 3). The gradual increase of stimulus intensity is a standard electrophysiological technique to quantify the number of presynaptic inputs innervating a postsynaptic neuron.

**Figure 3.**
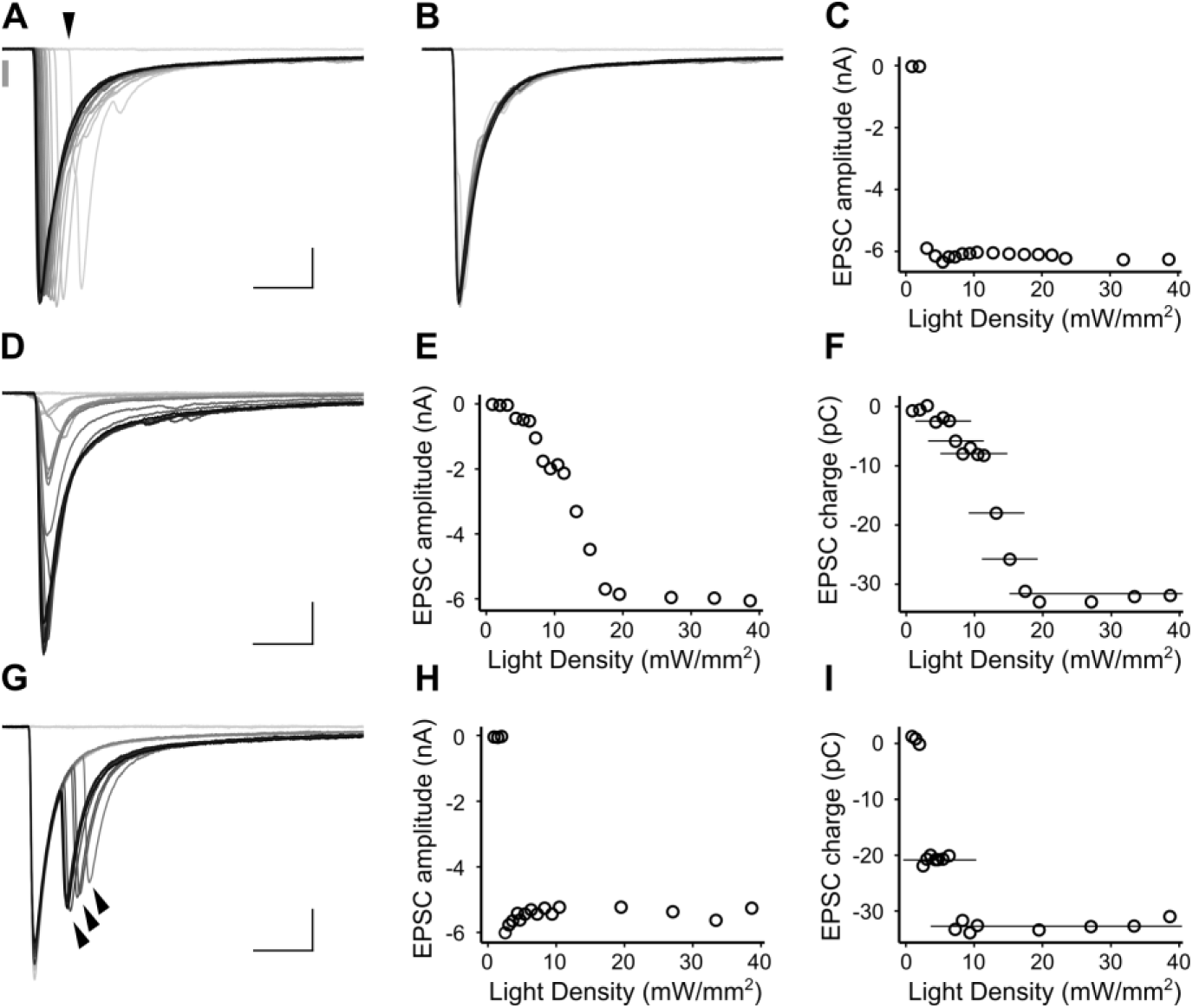
Gradual increase of light density evoked discrete EPSC steps in VL neurons. VL neurons in control mice were photostimulated by gradually increased light density, and the evoked currents were overlaid. The gray rectangles indicate the timing of photostimulation. Each current trace is color-coded; a darker gray represents a higher light density. Scale bars: 1 nA and 5 msec. **(A)** An example of all-or-none EPSCs. The arrowhead indicates the onset of the EPSC evoked by the minimum light density above the threshold. Note that the EPSC onsets shifted earlier as the light density increased. **(B)** The EPSC traces in panel A are aligned with the EPSC onsets. **(C)** A scatter plot showing the relationship between the light density and the EPSC amplitude. **(D-I)** Representative neurons showing multiple discrete EPSC steps. Panels D-F represent one neuron, and Panels G-I represent another neuron. The EPSC traces are aligned with the EPSC onsets (D and G). Scatter plots show the relationship between the light density and the EPSC amplitude (E and H) or charge transfer (F and I). The horizontal lines in panes F and I indicate the discrete EPSC steps. Note that the neuron in panel G initially showed one EPSC, but as light density increased, a delayed EPSC was evoked in addition to the first EPSC. The onset of the delayed EPSC shifted earlier as the light density further increased (arrowheads).

The gradual increase of light density, from 0 to ∼40 mW/mm^2^, yielded all-or-none EPSCs in some VL neurons, i.e., no EPSC was evoked when the light density was below a threshold, and evoked EPSCs essentially unchanged above the threshold (Fig 3A-C). The interval from photostimulation to EPSC onset always became shorter as the light density increased (Fig. 3A), presumably because stronger light activated more ChR2, causing faster depolarization of presynaptic terminals. When aligned with the onset, these EPSCs were identical (Fig. 3B), indicating that the recorded neuron was innervated by only one ChR2-expressing cerebellothalamic axon.

In other VL neurons, several discrete steps of EPSCs (Fig. 3D-F) or additional EPSCs with delayed onset (Fig. 3G-I) were evoked as the light density increased. The stepwise, non-continuous increase of EPSC amplitudes indicates that the recorded neuron was innervated by a relatively small number of ChR2-expressing cerebellothalamic axons; more of them were activated by stronger light (Fig. 3D-F). If the kinetics of ChR2-mediated presynaptic depolarization substantially differs among the axons, their EPSC peaks may not align, as shown in Figure 3G. In these cases, the charge transfer of EPSCs represents the number of inputs more precisely than the EPSC amplitude (Fig. 3H and I). Therefore, we used the number of discrete steps in the EPSC charge transfer as the number of ChR2-expressing cerebellothalamic axons innervating to the recorded neurons (Fig. 3F and I).

Since we cannot stimulate uninfected neurons using light, we likely underestimated the number of inputs in some cells. However, even if a neuron was innervated by only one or two ChR2-expressing cerebellothalamic axons (Fig. 3A and G), large EPSCs (∼ -6 nA) could be evoked, which were comparable to a neuron innervated by six ChR2-expressing cerebellothalamic axons (Fig. 3D, the most EPSC steps in our data set). Therefore, a small number of EPSC steps does not mean insufficient stimulation of cerebellothalamic axons. While it is difficult to quantify the number of inputs per cell accurately, our data suggest that the VL neurons receive a relatively small number of cerebellar inputs, like the primary sensory thalamus receives the sensory driver inputs.

### Developmental effects of Tsc1 deletion on the formation of cerebellothalamic synapses

In the primary visual and somatosensory thalamus, perturbing neuronal activity alters the synaptic strength, the AMPA/NMDA ratio, and the pruning of redundant driver inputs (Hooks & Chen, 2006; Wang & Zhang, 2008; Takeuchi *et al*., 2014). Considering the similar synaptic organization between the VL and these sensory thalamic nuclei, we hypothesized that early cerebellar perturbation alters the synaptic properties in the VL. To test this hypothesis, we compared the Purkinje cell-specific mutant mice—*Tsc1* deletion mice and Purkinje cell ablation mice—with their littermate control.

For Purkinje cell-specific deletion of *Tsc1*, we used the single transgenic mice (*Pcp2^Cre/+^* or *Tsc1^flox/+^*) as a control group and compared them with their double transgenic littermates for hemizygous deletion (*Pcp2^Cre/+^ Tsc1^flox/+^*). Previous studies showed that hemizygous deletion of *Tsc1* reduces the firing rate of Purkinje cells, and the mutant animals exhibit impaired social behaviors and associative sensory learning (Tsai *et al*., 2012, 2018; Kloth *et al*., 2015; Gibson *et al*., 2022). Homozygous deletion causes more robust phenotypes, but it also results in a substantial loss of Purkinje cells (Tsai *et al*., 2012, 2018; Gibson *et al*., 2022). To differentiate the potential effects of *Tsc1* deletion from Purkinje cell loss, we only used the hemizygous deletion model in this study.

The number of discrete EPSC steps was 3.00 ± 1.13 (n = 15 cells) in the control mice and 3.00 ± 0.97 (n = 16 cells) in the *Tsc1* deletion mice (TSC mice), indicating that each VL neuron receives a similar number of cerebellothalamic inputs regardless of *Tsc1* deletion (Fig. 4A-C). Other synaptic properties, i.e., the maximum amplitude and charge transfer of AMPA-EPSCs (Fig. 4E and F), the maximum amplitude of NMDA currents (NMDA-EPSCs, Fig. 4G), the AMPA/NMDA ratio (Fig. 4H), the 10-90% rise time and the weighted decay constant of AMPA-EPSCs (Fig. 4I and J, see Methods), are summarized in Table 1. The observed mean difference between the control and TSC mice and its bootstrap resampling distribution are shown as “TSC *minus* Control” in Figures 4C and E-J. No statistically significant difference was observed in any of these synaptic properties, as the 95% confidence interval of the estimated mean difference (thick vertical lines in Fig. 4C and E-J) included zero in all cases.

**Figure 4.**
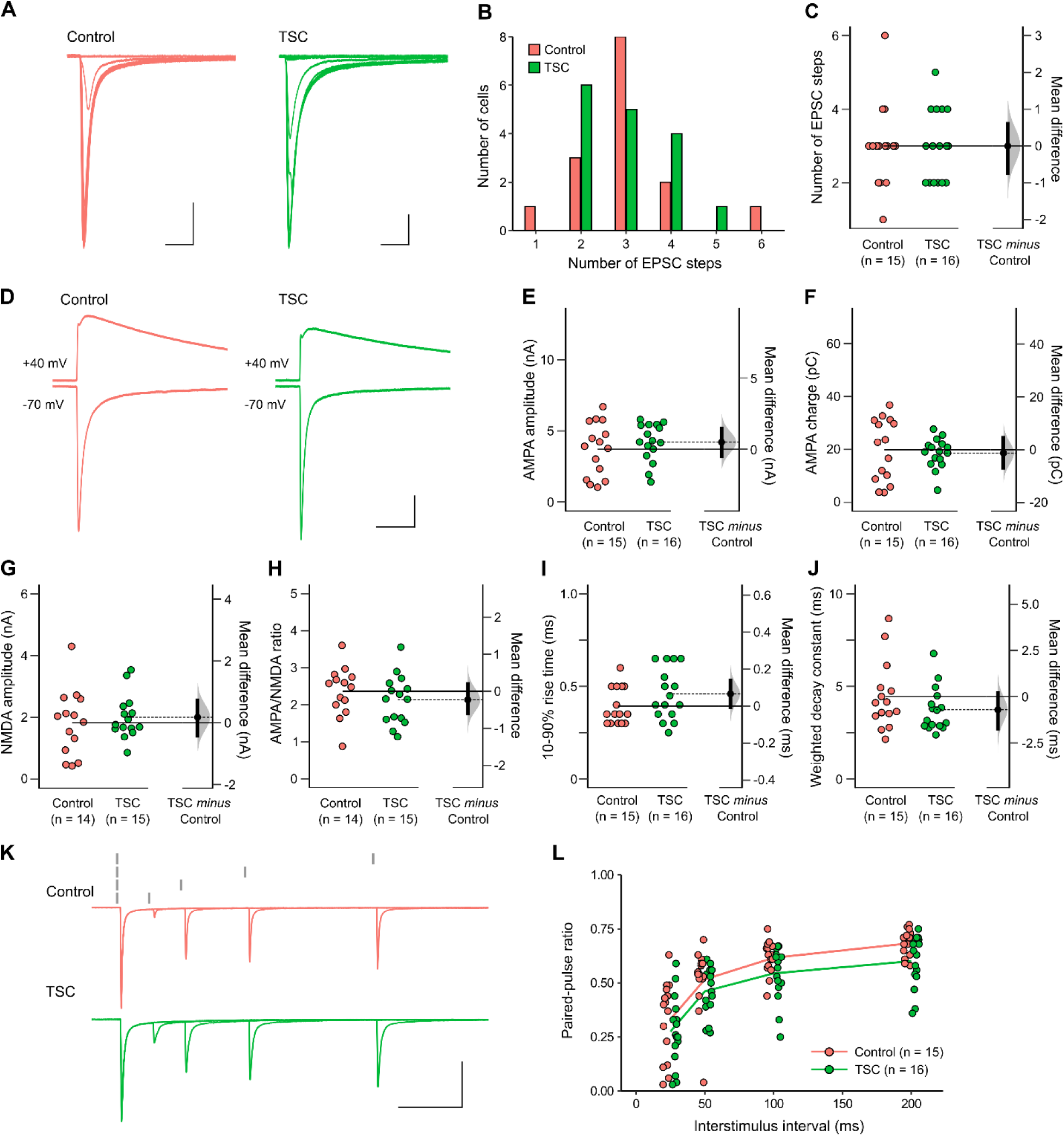
Physiological properties of cerebellothalamic synapses in *Tsc1* deletion mice. **(A)** AMPA-EPSCs evoked by gradually increased light density. Scale bars: 1 nA and 5 msec. **(B and C)** The number of EPSC steps was compared between the control and *Tsc1* deletion mice (TSC). The mean difference was estimated by bootstrap resampling. The gray curve in panel C shows the resampled distribution of the mean difference between the groups (TSC *minus* Control). The black circle and the thick vertical line indicate the observed mean difference and 95% confidence interval, respectively. The horizontal dashed line and solid line indicate the observed mean difference and zero mean difference, respectively. These lines are overlapped in this graph because the observed mean difference was zero. The statistical significance (*p* < 0.05) is assessed whether the 95% confidence interval includes the zero line. All the estimation graphics used in this study have the same structure. **(D)** EPSCs recorded from the same cells at -70 mV and +40 mV. Scale bars: 1 nA and 20 msec. **(E-J)** The amplitude (E) and charge transfer (F) of AMPA-EPSCs, the amplitude of NMDA-EPSCs (G), the AMPA/NMDA ratio (H), the 10-90% rise time (I), and the weighted decay constant (J) of AMPA-EPSCs were compared between the control and TSC mice. The statistical significance in the mean difference (TSC *minus* Control) is assessed as described above. Note that one cell in each group was lost before NMDA-EPSCs were recorded. It reduced the number of cells in the measurements requiring NMDA-EPSCs (G and H). **(K)** Paired-pulse depression with the interpulse interval of 25, 50, 100, and 200 msec. The gray rectangles above the EPSC traces indicate the timing and varying intervals between the two light pulses. Scale bars: 2 nA and 50 msec. **(L)** The paired-pulse ratio was compared between the groups.

**Table 1.** The summary of voltage-clamp recordings in the control and TSC mice.

|  | Control, n = 15<br>Mean (SD) | TSC, n = 16<br>Mean (SD) | 95% CI for the mean difference*<br>(TSC *minus* Control) |
| --- | --- | --- | --- |
| Series resistance (MΩ) | 7.26 (1.73) | 6.25 (1.45) | [-2.1, 0.0509] |
| Number of EPSC steps | 3.00 (1.13) | 3.00 (0.97) | [-0.788, 0.654] |
| AMPA amplitude (nA) | 3.72 (1.88) | 4.21 (1.34) | [-0.615, 1.59] |
| AMPA charge (pC) | 19.86 (11.71) | 18.56 (5.63) | [-7.51, 5.18] |
| NMDA amplitude (nA) | 1.83 (1.07) | 2.00 (0.71) | [-0.481, 0.789] |
| AMPA/NMDA ratio | 2.36 (0.66) | 2.14 (0.66) | [-0.649, 0.249] |
| 10-90% rise time (ms) | 0.39 (0.10) | 0.46 (0.14) | [-0.0157, 0.149] |
| Weighted decay constant (ms) | 4.44 (1.82) | 3.74 (1.18) | [-1.85, 0.299] |

\* Bootstrap confidence interval (CI) by 5000 resampling

We also quantified the paired-pulse ratio of AMPA-EPSCs, which is primarily determined by the presynaptic release probability (Fig. 4K and L). While the ratio was significantly affected by the interstimulus intervals (Mixed ANOVA, *F*(1.57, 45.66) = 133.27, *p* < 0.001), there was no significant difference between the control and TSC mice (*F*(1, 29) = 2.75, *p* = 0.11). Furthermore, no significant interaction was found between the mouse groups and the interstimulus intervals, i.e., the interstimulus intervals affected the paired-pulse ratio similarly in both groups of mice (*F*(1.57, 45.66) = 0.27, *p* = 0.71). These results suggest that hemizygous deletion of *Tsc1* does not alter either pre- or postsynaptic properties of cerebellothalamic synapses.

### Developmental effects of Purkinje cell ablation on the formation of cerebellothalamic synapses

Purkinje cells tonically inhibit the deep cerebellar nuclei, the source of cerebellothalamic axons. Thus, the ablation of Purkinje cells should substantially perturb the synaptic activity the thalamus receives from the cerebellum. To study the effects of Purkinje cell ablation, we used the single transgenic mice (*Pcp2^Cre/+^* or *Eno2^fsDTA/+^*) as a control group and compared them with their double transgenic littermates (*Pcp2^Cre/+^ Eno2^fsDTA/+^*) that lose most Purkinje cells within the first postnatal month (Fig. 1). The double transgenic mice (DTA mice) are slightly lighter than the control littermates and show moderate ataxia. However, despite the substantial cerebellar atrophy, they were healthy and showed no signs of health decline until we sacrificed them for electrophysiological analysis.

The electrophysiological properties of cerebellothalamic synapses in the control and DTA mice are shown in Figure 5 and Table 2. The AMPA/NMDA ratio (Fig. 5H) and the kinetics of AMPA-EPSCs (Fig. 5I and J) were not affected by Purkinje cell ablation. The paired-pulse ratio was not affected either (Fig. 5K and L), as there was no significant difference between the control and DTA mice (Mixed ANOVA, *F*(1, 35) = 1.07, *p* = 0.31) or no significant interaction between the mouse groups and the interstimulus intervals (*F*(1.23, 43.13) = 0.75, *p* = 0.42).

**Figure 5.**
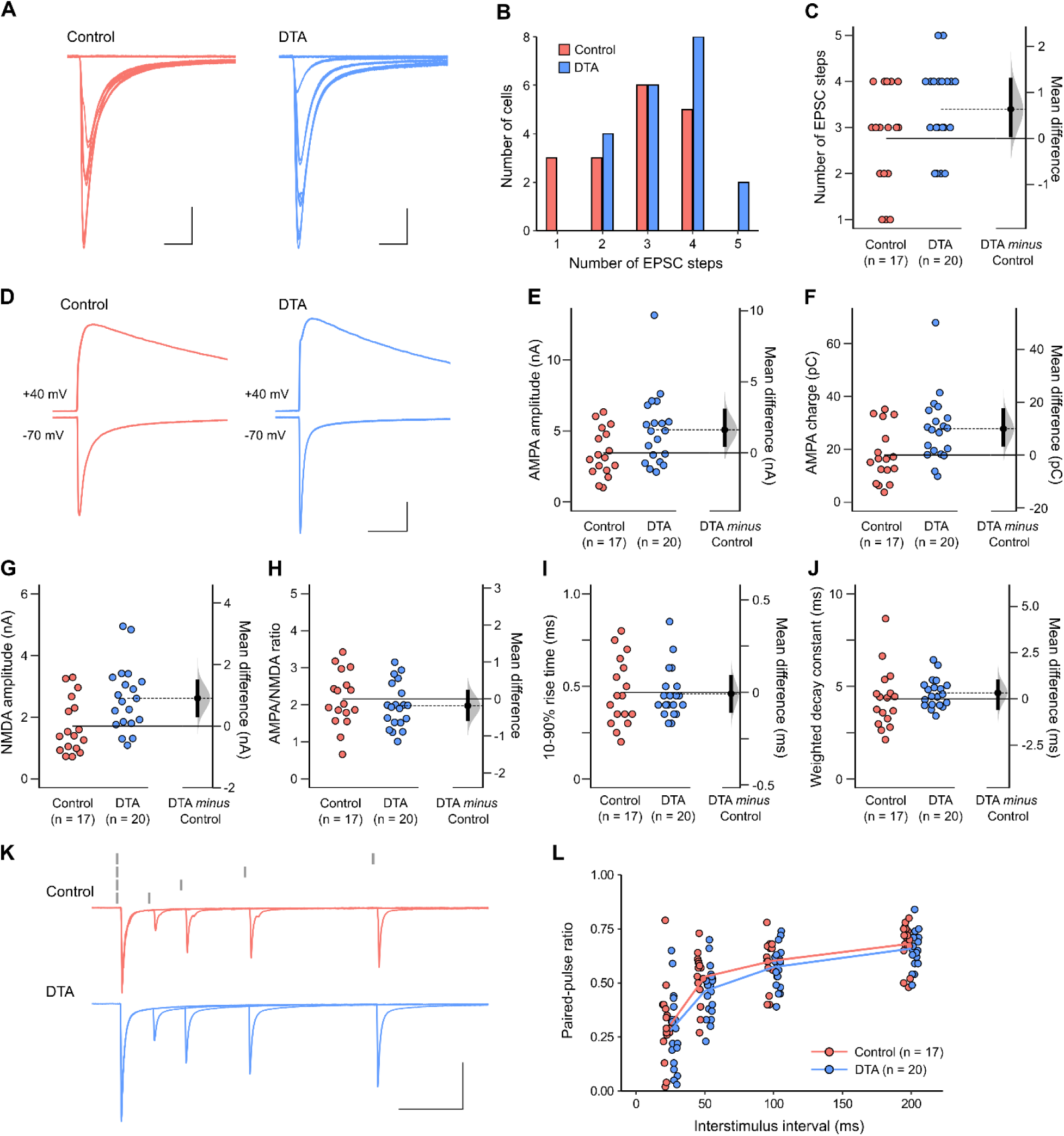
Physiological properties of cerebellothalamic synapses in PC ablation mice. **(A)** AMPA-EPSCs evoked by gradually increased light density. Scale bars: 1 nA and 5 msec. **(B and C)** The number of EPSC steps was compared between the control and the PC ablation mice (DTA). The mean difference was estimated by bootstrap resampling (DTA *minus* Control). The statistical significance was assessed as described in Figure 4. **(D)** EPSCs recorded from the same cells at -70 mV and +40 mV. Scale bars: 1 nA and 20 msec. **(E-J)** The amplitude (E) and charge transfer (F) of AMPA-EPSCs, the amplitude of NMDA-EPSCs (G), the AMPA/NMDA ratio (H), the 10-90% rise time (I), and the weighted decay constant (J) of AMPA-EPSCs. **(K)** Paired-pulse depression with the interpulse interval of 25, 50, 100, and 200 msec. The gray rectangles above the EPSC traces indicate the timing and varying intervals between the two light pulses. Scale bars: 2 nA and 50 msec. **(L)** The paired-pulse ratio was compared between the groups.

**Table 2.** The summary of voltage-clamp recordings in the control and DTA mice.

|  | Control, n = 17<br>Mean (SD) | DTA, n = 20<br>Mean (SD) | 95% CI for the mean difference*<br>(DTA *minus* Control) |
| --- | --- | --- | --- |
| Series resistance (MΩ) | 6.19 (1.29) | 6.38 (1.71) | [-0.699, 1.24] |
| # of EPSC steps | 2.76 (1.09) | 3.40 (0.94) | [0.0247, 1.32] |
| AMPA amplitude (nA) | 3.45 (1.66) | 5.07 (2.56) | [0.428, 3.12] |
| AMPA charge (pC) | 17.84 (10.35) | 27.77 (12.72) | [3.21, 17.8] |
| NMDA amplitude (nA) | 1.71 (0.89) | 2.62 (1.06) | [0.292, 1.51] |
| AMPA/NMDA ratio | 2.16 (0.74) | 1.97 (0.61) | [-0.602, 0.252] |
| 10-90% rise time (ms) | 0.47 (0.18) | 0.46 (0.14) | [-0.11, 0.094] |
| Weighted decay constant (ms) | 4.32 (1.61) | 4.64 (0.78) | [-0.607, 1.05] |

\* Bootstrap confidence interval (CI) by 5000 resampling

Unlike TSC mice, however, the other synaptic properties in the DTA mice differed from the control mice. The number of discrete EPSC steps was 2.76 ± 1.09 (n = 17 cells) in the control mice and 3.40 ± 0.97 (n = 20 cells) in the DTA mice (Fig. 5A-C). The 95% confidence interval of the mean difference between the groups (DTA *minus* Control) did not include zero, indicating that the difference was statistically significant (Fig. 5C). It should be noted, however, that the observed mean difference (0.64) was slight compared to the broad distribution of the bootstrap resampling, and the 95% confidence interval contains near zero values (Fig. 5C). These data suggest that the actual difference could be subtle or uncertain. On the other hand, DTA mice showed significantly larger AMPA-EPSCs and NMDA-EPSCs than the control mice with narrow 95% confidence intervals relative to the mean differences (Fig. 5E-G). These results suggest that the loss of Purkinje cells in the developing cerebellum strengthens the cerebellothalamic synapses without affecting the AMPA/NMDA ratio or presynaptic release probability. This strengthening might be associated with more cerebellothalamic inputs on each VL neuron, but its contribution is subtle, if any.

### Developmental effects of Tsc1 deletion on the excitability of thalamic neurons

Since cerebellar activities are relayed to the cerebral cortex as suprathreshold activities of thalamic neurons, abnormal thalamic excitability impacts cerebello-thalamo-cortical synaptic transmission. We therefore determined whether early cerebellar perturbations alter intrinsic membrane properties and excitabilities of VL neurons.

Under whole-cell current clamp mode, each VL neuron was excited by two different methods: synaptic stimulation by optogenetics (Fig. 6A and B) and somatic injection of depolarizing current (Fig. 6F and G). Although both methods evoked action potentials (APs) in nearly all VL neurons, synaptic stimulation elicited steep excitatory postsynaptic potentials (EPSPs), often making the AP threshold detection unreliable (Fig. 6A inset). Therefore, we measured the AP threshold, AP amplitude, and AP half-width by somatic current injection (Fig. 6H-J).

**Figure 6.**
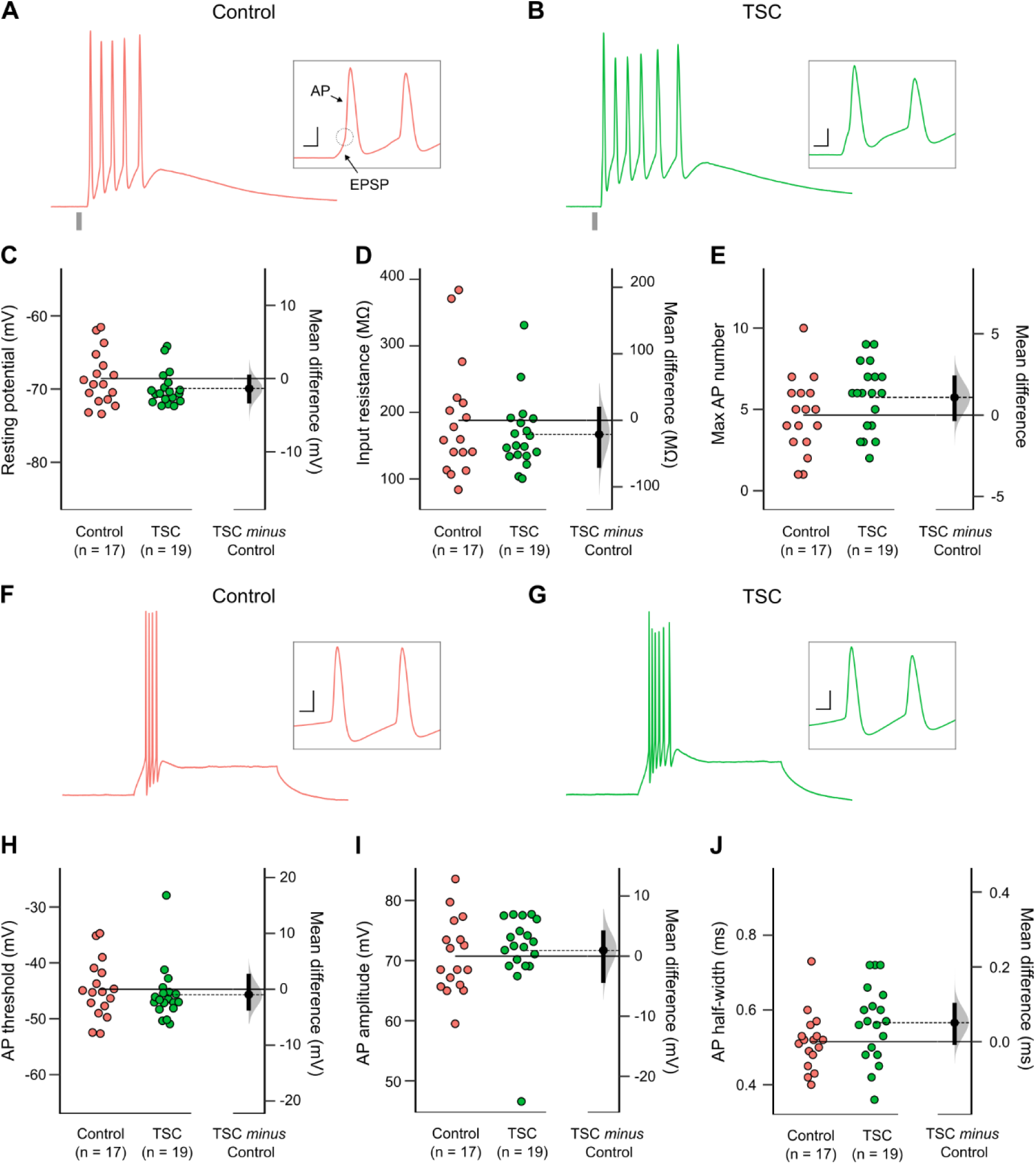
Membrane properties and excitability of VL neurons in *Tsc1* deletion mice. **(A and B)** APs evoked in VL neurons by photostimulation of cerebellothalamic axons in the control (A) and *Tsc1* deletion (TSC) mice (B). The gray rectangles indicate the timing of photostimulation. Insets: the first two APs are magnified along the time scale. Scale bars: 20 mV and 1 msec. Note that the first AP was evoked on the rising phase of EPSP, making the AP threshold detection unreliable (dashed circle). **(C-E)** The resting membrane potential (C), the input resistance (D), and the maximum number of synaptically evoked APs (E) were compared between the control and TSC mice. The mean difference between the groups was estimated by bootstrap resampling (TSC *minus* Control). **(F and G)** APs evoked in VL neurons by somatic current injection in the control (F) and TSC mice (G). Insets: the first two APs are magnified along the time scale. Scale bars: 20 mV and 1 msec. **(H-J)** The AP threshold (H), amplitude (I), and half-width (J) were compared between the control and TSC mice. The statistical significance in the mean difference (TSC *minus* Control) is assessed as described in Figure 4.

The resting membrane potential was measured in both methods before synaptic stimulation or current injection; then, the two values were averaged (Fig. 6C). The input resistance was measured by injecting hyperpolarizing current (Fig. 6D). The maximum number of synaptically evoked APs was measured by optogenetic stimulation using the highest light density (∼40 mW/mm^2^). As shown in Figure 3, this light density always evoked maximum EPSCs in our experimental condition, ensuring that all ChR2-expressing cerebellothalamic inputs were stimulated.

Figure 6 and Table 3 summarize these measurements in the control and TSC mice. Considering the normal organization of cerebellothalamic synapses in the TSC mice (Fig. 4), *Tsc1* deletion is unlikely to alter the membrane properties or excitabilities of VL neurons. It was indeed the case, as none of the measured values significantly differed between the control and TSC mice (Fig. 6).

**Table 3.** The summary of current-clamp recordings in the control and TSC mice.

|  | Control, n = 17<br>Mean (SD) | TSC, n = 19<br>Mean (SD) | 95% CI for the mean difference*<br>(TSC *minus* Control) |
| --- | --- | --- | --- |
| Resting potential (mV) | -68.57 (3.68) | -69.93 (2.32) | [-3.4, 0.544] |
| Input resistance (MΩ) | 188.00 (86.25) | 166.85 (53.79) | [-71.6, 20.6] |
| Max AP number | 4.65 (2.32) | 5.74 (2.10) | [-0.368, 2.44] |
| AP threshold (mV) | -44.73 (5.20) | -45.68 (4.93) | [-3.81, 2.77] |
| AP amplitude (mV) | 70.74 (6.18) | 71.7 (6.93) | [-4.44, 4.3] |
| AP half-width (ms) | 0.52 (0.08) | 0.57 (0.10) | [-0.00867, 0.104] |

\* Bootstrap confidence interval (CI) by 5000 resampling

### Developmental effects of Purkinje cell ablation on the excitability of thalamic neurons

The loss of Purkinje cells increased the charge transfer of AMPA-EPSCs by 55.7 % (control vs. DTA = 17.84 pC vs. 27.77 pC, Table 2). Whether this increase leads to enhanced thalamic firing depends on VL neurons’ intrinsic membrane properties and excitabilities.

The resting membrane potential of VL neurons was -69.45 ± 3.96 mV (n = 20 cells) in the control mice and -71.89 ± 3.45 mV (n = 16 cells) in the DTA mice (Table 4). The mean difference between the groups was slight (2.44 mV) yet statistically significant, as the 95% confidence interval narrowly excluded zero (Fig. 7C). These data indicate that although the difference between the groups is too subtle to be conclusive, VL neurons in the DTA mice might be slightly hyperpolarized than the control mice.

**Figure 7.**
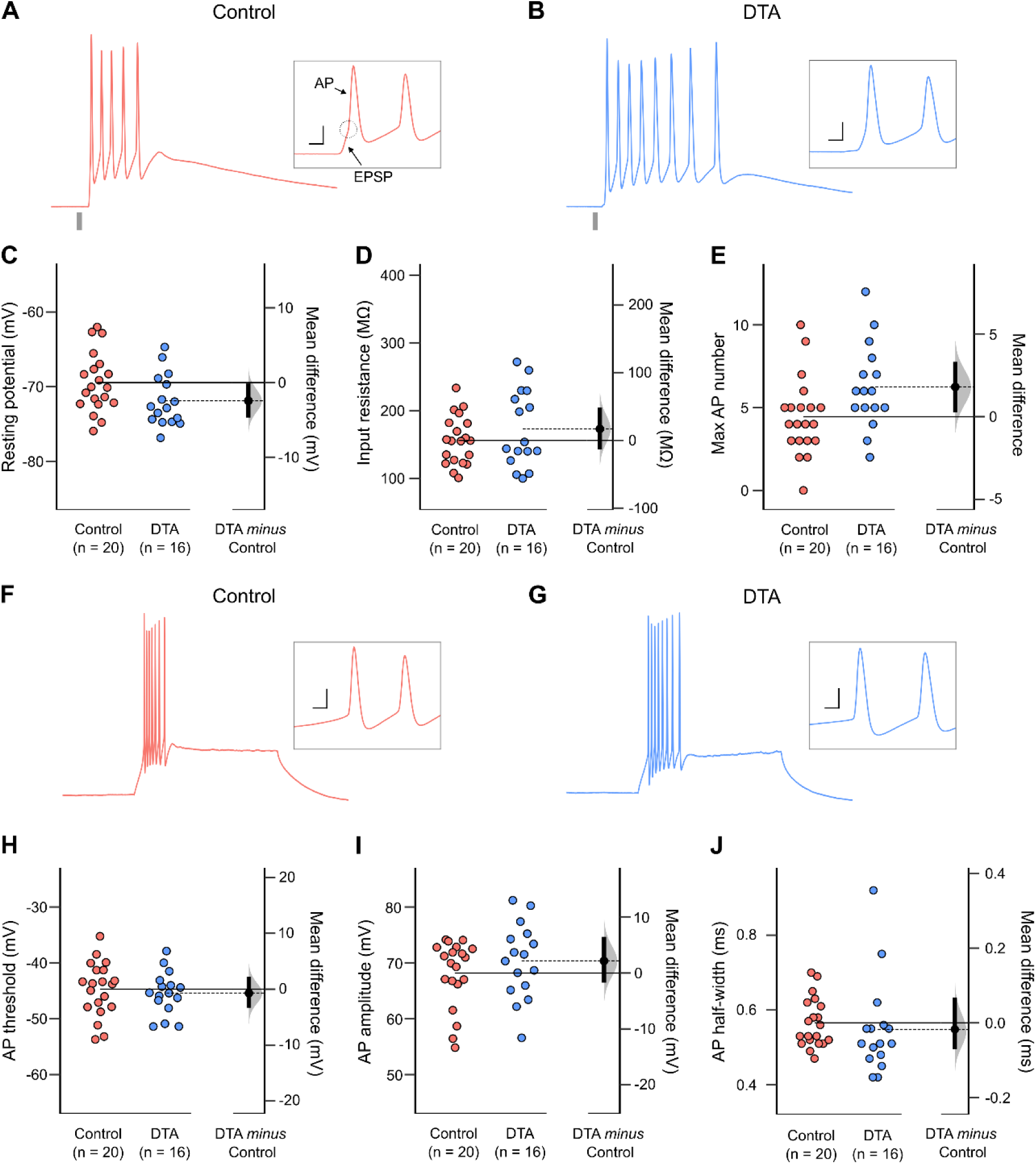
Membrane properties and excitability of VL neurons in PC ablation mice. **(A and B)** APs evoked in VL neurons by photostimulation of cerebellothalamic axons in the control (A) and PC ablation (DTA) mice (B). The gray rectangles indicate the timing of photostimulation. Insets: the first two APs are magnified along the time scale. Scale bars: 20 mV and 1 msec. Note that the first AP was evoked on the rising phase of EPSP, making the AP threshold detection unreliable (dashed circle). **(C-E)** The resting membrane potential (C), the input resistance (D), and the maximum number of synaptically evoked APs (E) were compared between the control and DTA mice. The mean difference between the groups was estimated by bootstrap resampling (DTA *minus* Control). **(F and G)** APs evoked in VL neurons by somatic current injection in the control (F) and TSC mice (G). Insets: the first two APs are magnified along the time scale. Scale bars: 20 mV and 1 msec. **(H-J)** The AP threshold (H), amplitude (I), and half-width (J) were compared between the control and DTA mice. The statistical significance in the mean difference (DTA *minus* Control) is assessed as described in Figure 4.

**Table 4.** The summary of current-clamp recordings in the control and DTA mice.

|  | Control, n = 20<br>Mean (SD) | DTA, n = 16<br>Mean (SD) | 95% CI for the mean difference*<br>(DTA *minus* Control) |
| --- | --- | --- | --- |
| Resting potential (mV) | -69.45 (3.96) | -71.89 (3.45) | [-4.7, -0.134] |
| Input resistance (MΩ) | 156.15 (35.79) | 172.97 (56.64) | [-13.3, 48.2] |
| Max AP number | 4.45 (2.33) | 6.25 (2.57) | [0.263, 3.31] |
| AP threshold (mV) | -44.71 (4.83) | -45.43 (3.86) | [-3.36, 2.24] |
| AP amplitude (mV) | 68.11 (5.97) | 70.37 (6.72) | [-1.64, 6.57] |
| AP half-width (ms) | 0.57 (0.07) | 0.55 (0.13) | [-0.0711, 0.0677] |

\* Bootstrap confidence interval (CI) by 5000 resampling

The input resistance, AP threshold, AP amplitude, and AP half-width were unaltered in the DTA mice, indicating that losing Purkinje cells does not change the subthreshold or suprathreshold excitability of VL neurons (Fig. 7D, H-J, and Table 4). However, previous studies showed that VL neurons discharge more APs when they are more hyperpolarized before being stimulated (Contreras & Steriade, 1995; Llinás & Steriade, 2006; Schäfer *et al*., 2021); thus, the resting membrane potential influences the AP firing pattern. The maximum number of synaptically-evoked APs was 4.45 ± 2.33 (n = 20 cells) in the control mice and 6.25 ± 2.57 (n = 16 cells) in the DTA mice (Table 4). The difference between the groups was statistically significant (Fig. 7E). It remains unclear whether the increased AP number in the DTA mice is due to increased AMPA currents (Fig. 5E and F) or the possible difference in the membrane potential (Fig. 7C). Nevertheless, these results suggest that losing Purkinje cells leads to enhanced inputs and outputs of VL neurons.

### The contribution of the resting membrane potential and synaptic strength to the synaptically-evoked action potentials in VL neurons

The membrane potential shortly before depolarization substantially affects how the VL neurons discharge APs (Contreras & Steriade, 1995; Llinás & Steriade, 2006; Schäfer *et al*., 2021). However, little is known about the extent to which thalamic AP firing depends on the strength of the cerebellothalamic inputs. To examine the contribution of synaptic strength to the AP number, we pooled all 37 control cells used to analyze the membrane properties and excitability of VL neurons (Fig. 6 and 7). Although these cells were recorded primarily under the current-clamp mode, voltage-clamp recordings were also performed to measure the size of the maximum AMPA-EPSCs. We excluded one cell that exhibited voltage-clamp failure. Using the remaining 36 control cells, we examined the factors that correlated with the maximum number of synaptically evoked APs.

The maximum number of synaptically evoked APs in individual VL neurons negatively correlated with their resting membrane potential (Fig. 8A, Pearson’s correlation coefficient, *r*(34) = -0.38, *p* = 0.021). This result is consistent with the previous studies, showing that VL neurons evoke more APs when they are more hyperpolarized before stimulation. Interestingly, the maximum AP number also positively correlated with the charge transfer of the maximum AMPA-EPSCs they received (Fig. 8B, r(34) = 0.48, *p* = 0.003).

**Figure 8.**
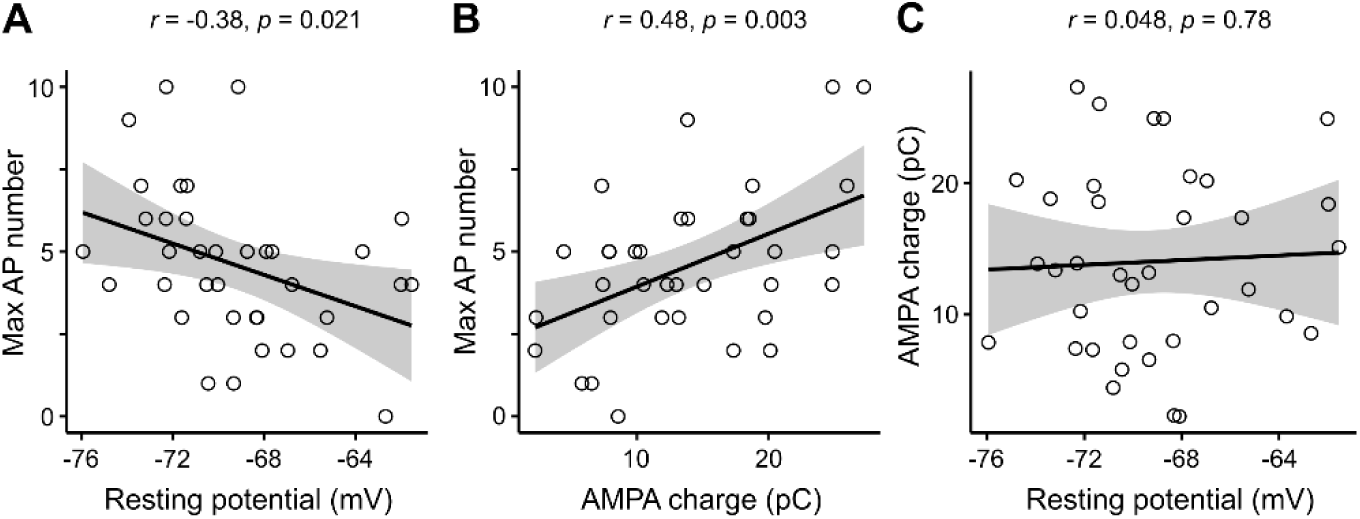
Factors affecting synaptic outputs of VL neurons. (**A**) Correlation between individual neurons’ resting membrane potential and the maximum number of synaptically evoked APs. (**B**) Correlation between individual neurons’ synaptic charge transfer by the maximum AMPA-EPSCs and the maximum number of synaptically evoked APs. (**C**) Correlation between individual neurons’ resting membrane potential and synaptic charge transfer by the maximum AMPA-EPSCs. In all plots, the solid lines indicate linear regression lines, and the gray shades indicate their 95% confidence intervals.

If VL neurons receiving stronger synaptic inputs were coincidentally more hyperpolarized in our data set, the maximum AP number mistakenly appears to correlate with the synaptic strength. To test this unlikely scenario, we examined the correlation between the resting membrane potential and the charge transfer of the maximum AMPA-EPSCs. The correlation coefficient was near zero, showing that these two values were uncorrelated (Fig. 8C, r(34) = 0.048, *p* = 0.78). These results indicate that stronger cerebellothalamic inputs can evoke more APs in VL neurons. Considering the subtle, uncertain difference in the resting membrane potential between the control and DTA mice, we assume that VL neurons in the DTA mice exhibited more synaptically-evoked APs primarily because of their enhanced synaptic inputs from the cerebellum.

## Discussion

In this study, we examined how cerebellar perturbation early in life affects the formation of cerebellothlamic circuits in the VL. The VL is the primary thalamic target of the cerebellum in rodents, making it the ideal model for gaining basic biological insight into how the cerebellum affects thalamic development. We found that a relatively small number of cerebellothalamic axons provide a powerful excitatory drive to individual VL neurons in control mice. This synaptic organization is similar to sensory inputs innervating the primary sensory thalamus, which is known to be refined by neuronal activity during development. We also found that the loss of Purkinje cells enhanced synaptic inputs and outputs of VL neurons, suggesting that the activity of the deep cerebellar nuclei tunes the development of cerebellothalamic synapses. Since the organization of the cerebello-thalamo-cortical circuit is essentially the same between motor and non-motor systems, what we learn from the VL may also be applicable to non-motor systems.

### The number of cerebellothalamic inputs per VL neuron

Our data showed that individual VL neurons receive 1-6 cerebellothalamic axons, approximately 3 on average. This number is likely fewer than the actual number of inputs because not all the deep cerebellar nuclei neurons expressed ChR2. However, electrical stimulation is not a better alternative because it is non-selective, and a stimulation electrode must be placed 0.5-1 mm away from the recording site to avoid stimulating corticothalamic axons (Aumann *et al*., 2000). In this configuration, axons that run at an oblique angle to the tissue surface are prone to be severed by slicing and difficult to excite.

Previous studies used electrical stimulation to quantify the number of sensory driver inputs innervating individual neurons in the primary visual and somatosensory thalamus. These studies reported 1-3 sensory driver inputs, mostly one input per neuron (Chen & Regehr, 2000; Arsenault & Zhang, 2006; Hooks & Chen, 2006; Wang & Zhang, 2008; Takeuchi *et al*., 2014). However, a recent study showed that optogenetically evoked EPSCs are significantly larger than electrically evoked EPSCs in the primary visual thalamus, indicating that electrical stimulation indeed fails to excite some inputs. It is currently estimated that approximately 10 retinal ganglion cell axons innervate a single neuron in the primary visual thalamus, among which around 3 inputs are strong enough to drive postsynaptic AP firing (Litvina & Chen, 2017).

The number we reported herein was fewer than the above estimate, but many VL neurons were innervated by 2 strong inputs, i.e., an input larger than 1 nA amplitude that almost always evokes thalamic APs. Therefore, we assume that the number of driver inputs per neuron is similar between the VL and the primary visual thalamus.

It remains unclear whether the cerebellothalamic synapses undergo synapse elimination. i.e., developmental decline of the number of inputs per neuron, as the primary visual and somatosensory thalamus does. AAV needs 2-3 weeks until the gene expression reaches a sufficient level; thus, it is unsuitable for performing optogenetics in neonatal animals. The main goal of this study was not to determine whether VL neurons initially receive more inputs from the cerebellum. Instead, we aimed to reveal the consequences of early cerebellar perturbation on the formation of thalamic circuits. Our data showed that, regardless of whether synapse elimination occurs, neither *Tsc1* deletion nor Purkinje cell ablation substantially changes the number of cerebellar inputs per neuron in mature VL.

### Hemizygous deletion of Tsc1 in Purkinje cells

Although Purkinje cells spontaneously discharge action potentials, their firing rate is modulated by excitatory and inhibitory synaptic inputs. Inhibitory inputs are provided by interneurons in the cerebellar molecular layer (MLI), and chemogenetic excitation and inhibition of MLI both lead to impaired social behavior in mice (Badura *et al*., 2018; Chao *et al*., 2023). These results suggest that aberrant Purkinje cell activity is not a coincidence but a cause of social and cognitive impairments. To test whether aberrant Purkinje cell activity also affects the development of the cerebellothalamic circuit, we used the Purkinje cell-specific *Tsc1* deletion model. Since *Tsc1* deletion reduces the firing rate of Purkinje cells, it is reasonable to hypothesize that Purkinje cell hypoactivity upregulates the activity of the deep cerebellar nuclei, affecting the development of cerebellar target regions. However, synaptic properties and excitability of the cerebellothalamic circuit were not affected by *Tsc1* deletion. Although there remains a possibility that the motor and non-motor thalamus have different susceptibilities to aberrant cerebellar activities, our data suggest that moderate changes in Purkinje cell firing rate do not affect thalamic development.

### Purkinje cell ablation

Purkinje cell abnormalities are commonly associated with neurological conditions. The gene expression profile in the prenatal human cerebellum revealed that Purkinje cells express more than 50 ASDs related genes involved in neuronal development, suggesting the crucial roles of Purkinje cells in developing motor and non-motor circuits (Sydnor & Aldinger, 2022). Furthermore, a previous study using a spontaneous mutant mouse, Lurcher, showed that the degree of Purkinje cell loss significantly correlated with the degree of repetitive behaviors and hyperactivity (Martin *et al*., 2010).

While Purkinje cell ablation caused substantial cerebellar atrophy (Fig. 1), we did not notice any decrease in the tdTomato-expressing cerebellothalamic axon terminals in the VL. It indicates that most deep cerebellar nuclei neurons survived despite the considerable cerebellar malformation after the AAV injection. However, unlike *Tsc1* deletion, Purkinje cell ablation strengthens cerebello-thalamo-cortical synaptic transmission. It suggests that the maturation of the cerebellothalamic circuit depends on the cerebellar activity to some extent and that abnormal upregulation of deep cerebellar nuclei—due to Purkinje cell loss—enhances the synaptic strength.

In addition, the resting membrane potential of the VL neurons tended to be slightly hyperpolarized by Purkinje cell ablation. This result was statistically significant yet uncertain because the mean difference between the control and Purkinje cell ablation mice was small relative to the variation within a group. Still, a homeostatic mechanism might hyperpolarize the VL neurons in the Purkinje cell ablation mice to compensate for their enhanced synaptic inputs. It should be noted that even a few millivolts more hyperpolarization before excitation results in more action potentials in VL neurons. Therefore, stronger synaptic inputs and more hyperpolarized postsynaptic membranes both enhance the outputs of VL neurons (Fig. 8).

### Enhancement vs. suppression of cerebellothalamic activity

Compared to the previous studies on the primary sensory thalamus, neuronal activity had relatively minor effects on thalamic development in the VL. One caveat is that the activity of sensory driver inputs was blocked in the previous studies, whereas Purkinje cell reduction (DTA mice) or hypoactivity (TSC mice) enhances cerebellothalamic activity. Although these conditions mimic common cerebellar abnormalities in human patients and animal models of neurological disorders, reducing cerebellothalamic activity may impact the development of the cerebellothalamic circuit more. In this regard, it is interesting to study cerebellothalamic synapses in disease models associated with Purkinje cell hyperactivity, such as fetal alcohol syndrome and the gene mutation associated with amyotrophic lateral sclerosis and frontotemporal dementia (Servais *et al*., 2007; Liu *et al*., 2022).

In human patients with developmental cerebellar lesions, isolated cerebellar damage reduces the volume of the contralateral cerebral cortex (Limperopoulos *et al*., 2010, 2014; Stoodley & Limperopoulos, 2016). This phenomenon is unlikely to be explained by the upregulation of cerebellothalamic activities. Instead, it is more reasonable to assume that damage to the deep cerebellar nuclei causes hypoactivity or degeneration of the cerebellothalamic axons, reducing synaptic activity in the cerebral cortex. Indeed, magnetic resonance imaging revealed that cerebellothalamic axons seem somewhat damaged in some patients after pediatric tumor surgery (Morris *et al*., 2009; Law *et al*., 2011). When one side of cerebellothalamic axons is transected in neonatal animals, new collaterals sprout from the spared side and reinnervate the deafferented thalamus (Molinari *et al*., 1986). A similar circuit reorganization might occur in pediatric cerebellar surgery in human patients and contribute to the pathogenesis of motor and non-motor impairments.

In summary, early cerebellar damage affects thalamic development depending on the type and severity of damage. Although its functional significance in neurological disorders is yet to be elucidated, abnormal thalamic development likely alters how the cerebellum regulates motor and non-motor behaviors throughout life.

## Acknowledgments

We thank Daniel Johnston, Richard Gray, and Darrin Brager for their helpful support in conducting electrophysiology experiments and Shigeyoshi Itohara for kindly providing us with the *Eno2^fsDTA/+^* transgenic mice.

